# The transmission of Hand, Foot, and Mouth Disease in East and Southeast Asia

**DOI:** 10.1101/612580

**Authors:** Jijun Zhao, Yanfen Wang

## Abstract

Hand Foot and Mouth Disease (HFMD) is in endemic in many countries in East and Southeast Asia, including those in the tropical or subtropical climate zones. To substantially reduce HFMD, it is necessary to design effective control measures, which is based on a deep understanding of the disease transmission. However, the transmission mechanism of HFMD was rarely studied. The cyclic pattern of HFMD incidence is believed to be related to climatic factors, rather than school terms as observed from childhood infectious diseases in developed countries in the prevaccination era. Furthermore, the association of incidence and climatic factors in different locales in China are inconsistent and even contradictory. Here we selected countries or regions in typical climatic zones in East and Southeast Asia to study the transmission rate and its seasonality for HFMD. Countries or regions selected representing temperate, subtropical and tropical zones are Japan, Hong Kong SAR, Macau SAR and Singapore. Comparatively, we chose provinces in mainland China in three climate zones and contrast them with above selected regions or countries. We used Time Series Susceptible Infected Recovered (TSIR) model to estimate the HFMD transmission rate. The parameters in the TSIR model were estimated by Markov Chain Monte Carlo (MCMC). We then used a linear regression model to analyze the effects of climate factors, seasonal contact rate in children (and seasonal contact rate in population for provinces in China) on the transmission rate of HFMD in selected regions. We found that: 1) transmission rate of HFMD is highly seasonal in the studied countries, SARs and provinces of mainland China, except Singapore; 2) the HFMD transmission rate can be affected by the climatic factors as well as the seasonal contact rate of population, depending on which factor is dominant; 3) The transmission rate in provinces in China increased dramatically during the time period of Chinese Spring Travel Rush that has higher population contact; 4) transmission rate seasonality in Japan, Hong Kong SAR and Macau SAR is affected by climatic factors.

**Author Summary:** Hand, Foot and Mouth Disease (HFME) is endemic in East and Southeast Asia with reported cases of more than two million every year. The epidemic patterns such as annual cyclic pattern of reported HFMD cases have been observed and studied for the purpose of understanding the disease. The mechanisms that describe how a disease is transmitted cannot be observed, however they lead to the observed epidemic patterns of the disease. We analyzed the transmission rate (that help to understand the transmission mechanism) of HFMD in selected countries or regions that represent territories in tropical, subtropical and temperate climatic zones in East and Southeast Asia and compared the HFMD transmission in these regions. We also analyzed the possible driving factors of the seasonal transmission of HFMD. The transmission of HFMD can be affected by both social behavior and climatic factors, however either of them can dominant the effect on HFMD transmission depending on regions or countries. In mainland China, high population contact rate is the dominant factor to have high HFMD transmission; while in Japan, Hong Kong SAR and Macau SAR, climatic factors have the dominant effect. These findings can help design effective control measures.

## Introduction

Hand, foot and mouth disease (HFMD) is a common infectious disease caused by a variety of enteroviruses, mainly coxsackievirus 16 (CVA16) and enterovirus 71 (EV71) [1]. Most of the patients are children under five years old [1,2]. The symptom of HFMD is usually mild and self-limiting, but the infection has also been rarely associated with neurological diseases and can even lead to death [3]. HFMD is now in endemic in many regions or countries in East and Southeast Asia, including those in the tropical or subtropical climate zones [4]. Because the Asia-Pacific region has the most HFMD incidence, HFMD is a serious threat to the health of children in countries in the region [5]. For childhood infectious diseases such as HFMD, a high-level-coverage vaccination is the effective control method. Started from late 2016, vaccination of enterovirus EV71 was put into voluntary use in various provinces in China [6], but there is no vaccination for HFMD in other regions or countries. To substantially reduce or eliminate an infectious disease, it is necessary to design and assess effectiveness of [control measures of the disease, which is based on a deep understanding of the disease transmission [5].

The transmissions of some childhood infectious diseases (including measles, rubella, pertussis, etc.) in developed countries in Europe and the United States have been intensively studied in the past few decades [7–10]. One of the important findings is that the transmission rate baseline, transmission rate seasonality and the susceptible pool determine the outbreak size of an infectious disease and the incidence periodicity. It is further found that the periodic contact rate of school aged children due to school terms lead to the transmission rate seasonality of some childhood infectious diseases. These findings not only help to explain the temporal and spatial patterns of observed incidence, but also help to develop cost-effective control measures. However, the transmission rate seasonality of HFMD has rarely been studied [11].

**Figure 1.**
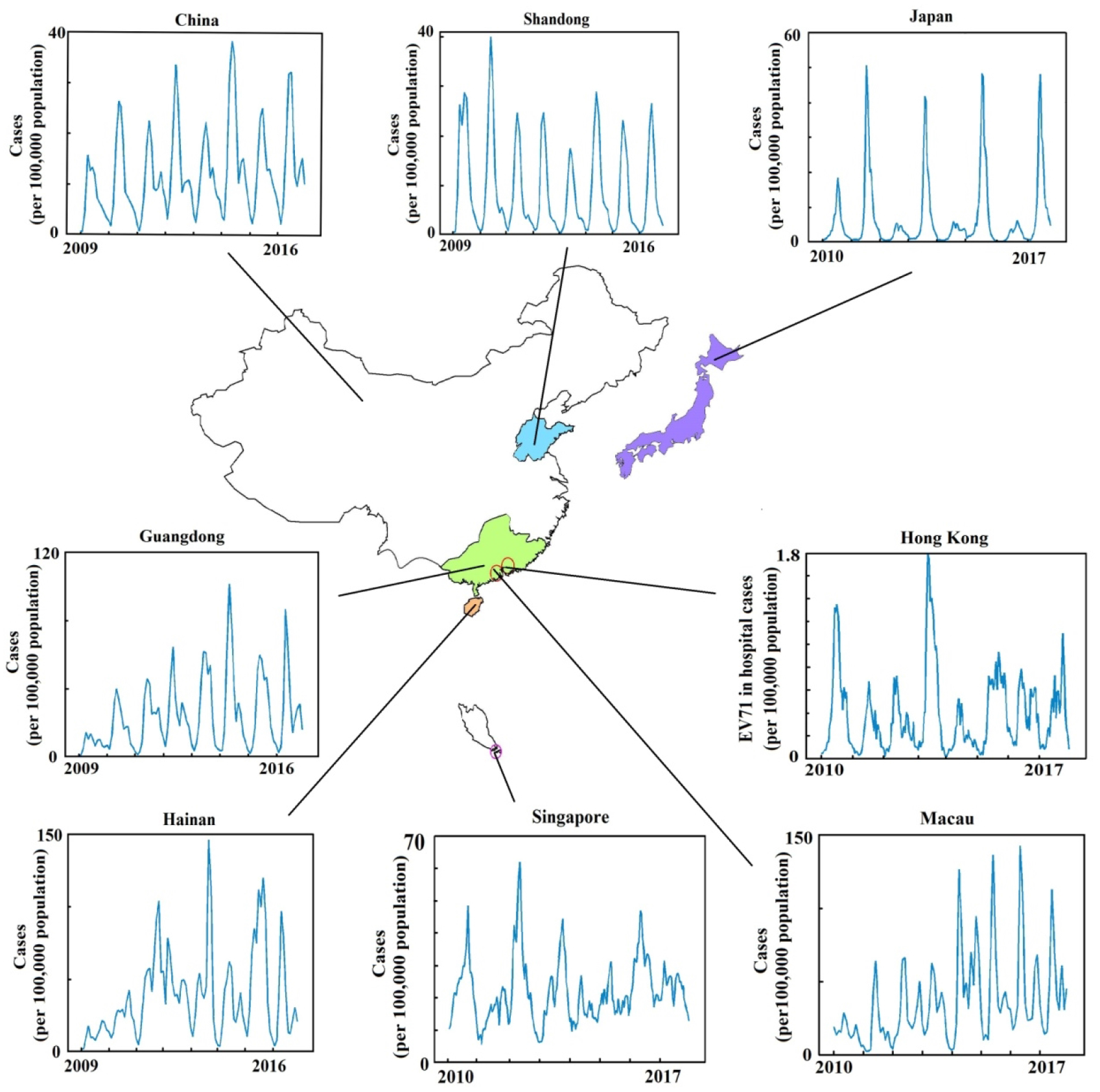
HFMD incidence patterns in Japan, Hong Kong SAR, Macau SAR, Singapore, and comparative provinces (Shandong, Guangdong, Hainan) in mainland China. This figure only shows maps of mainland China, Japan, Hong Kong SAR, Macau SAR, and Singapore, and does not include other regions or countries in East and Southeast Asia.

Similar to other childhood infectious diseases in developed countries in the prevaccination era, HFMD incidence in East Asia seems to have a cyclical pattern (Figure 1). However, the periodicity of HFMD incidence is believed to be related to climatic factors [12,13], rather than school terms for those childhood infectious diseases in developed countries. The associations of HFMD incidence with climatic factors in Vietnam, South Korea, Japan and Hong Kong are relatively consistent: with relatively higher temperature or humidity, higher incidences are expected [12–15]. Reviews of articles including many countries also found the significant association between HFMD cases and both temperature and relative humidity [16,17], and association between HFMD cases and more climatic factors [17]. However, the conclusions of the association of incidence cyclic pattern with climatic factors in different specific locales (province, city or county) in China are inconsistent and even contradictory [18–20]. Studies including many provinces in China only show a weak association between incidence and climatic factors [1]. Since an incidence cyclic pattern was determined by the transmission rate and the number of susceptible, to understand the transmission of HFMD, we need to study whether climatic factors affect the transmission rate and thus further affect the periodicity of the incidence. If the transmission rate is affected by climatic factors, then there might be a variation in the transmission rate seasonality of HFMD in different climatic zones. In the tropical zone, we may expect less amplitude of the seasonality of the transmission rate than in the temperate zone because some climatic factors change less in tropical area. China covers geographically tropical, subtropical, temperate, and other climatic zones, hence we may also see a variation in transmission rate in different provinces in China.

Here we select countries or regions in typical climatic zones to study the transmission rate and its seasonality for HFMD. We choose Singapore and Japan for the tropical and the temperate zones, and Hong Kong Special Administrative Region (SAR) and Macau SAR for the subtropical climate zone. We select these regions or countries to cover three typical climatic zones, where HFMD has been circulated in endemic, and we also consider whether these regions have data of consecutive several years or not such that transmission patterns can be estimated. Comparatively, we chose provinces in mainland China in three climate zones and contrast them with above selected regions or countries. Provinces selected in mainland China include Hainan, Guangdong and Shandong representing regions in tropical, subtropical and temperate zones in China. We will study the transmission rate and its seasonality of HFMD in different countries and regions, and to see whether there is any variation in the transmission rate seasonality.

We will further analyze the factors affecting the seasonality of HFMD transmission rate, including climatic factors and school terms. A factor that will be typically considered for mainland China is the seasonal population contact rate. The population contact rate can be affected by seasonal population aggregation and seasonal population flux. Mainland China has the largest seasonal population flux in the world due to the Chinese Spring Festival (CSF), which is called the Spring Festival Travel Rush (SFTR). Officially, the SFTR lasts 40 days every year, 15 days before and 25 days after the Lunar New Year. During SFTR, people travel back and forth from places of working to their hometowns. The closed and crowded carriages of long-distance buses and crowded population in waiting halls of long-distance bus stations and train stations provide good conditions for the spread of the disease. In addition, during the CSF, people visiting relatives and friends, hence the contact rate of the population increased.

Hence, besides climatic factors and school terms, the time frame of SFTR will also be considered in the influential factor on HFMD transmission rate for provinces in mainland China. We will study whether the transmission rate seasonality of HFMD in different countries or regions in East and Southeast Asia has a general driver or has different drivers. We try to explain why associations of climatic factors and incidence in different regions in mainland China have consistent or contradicted conclusions.

## The Data

### Reported cases of HFMD

The incidence data in Hainan, Guangdong and Shandong provinces and the entire mainland China were monthly reported data from 2009 to 2016 and was obtained from The Data Center of China Public Health Science [21]. Monthly reported cases of enterovirus infection in Macau SAR came from the Communicable Prevention and Diseases Surveillance Unit of the Health Bureau of Macau SAR [22].

Two types of HFMD data can be obtained from the Department of Health of the Hong Kong SAR [23]. One is monthly in-hospital cases of enterovirus EV71 (no cases of other types of viruses), another type is the consultation rate of general practitioners. Since consultation rates are not reported cases and cannot be utilized by our method, we use the in-hospital cases. From the official report of the Department of Health of Hong Kong SAR, HFMD caused by EV71 accounted for around 6% of the total detection. However, this will not affect our estimation of the transmission rate and the basic reproduction number of EV71 in Hong Kong because the reporting rate will be counted. We will see a much lower reporting rate in EV71 in Hong Kong.

The reported cases of HFMD in Singapore and Japan are weekly data, and we aggregate them into bi-week data and four-week data. Data for Singapore were obtained from the Ministry of Health of Singapore [24], and data for Japan were obtained from the National Institute of Infectious Diseases in Japan [25]. Data for Macau SAR, Hong Kong SAR, Singapore and Japan are from 2010 to 2017.

### Demographic data

The population and annual birth rate in Hainan, Guangdong and Shandong provinces and in the entire mainland China were obtained from the National Bureau of Statistics of China [26], and the number of births of each province is calculated according to the two. The population and births in Macau, Hong Kong, Singapore and Japan were obtained from their respective statistical departments [22,27,28]. The number of births in Japan is monthly data; population and births in Macau, Hong Kong and Singapore are yearly data.

### School terms and Spring Festival Travel Rush

School terms in Hong Kong SAR follow guidelines from the Education Bureau of Hong Kong with the convention that school starts on the 1st of September and ends in early July. Besides, there is a ten-day Christmas and New Year holiday, and a one-week CSF Holiday. Macau SAR has the similar school terms and vacations as Hong Kong SAR. We refer school terms in mainland China to announcements by provincial education departments. Summer vacation is usually in July and August, and winter vacation is usually four to five weeks in January and February depending on the lunar calendar date of CSF. In Japan, there are three vacations or breaks a year: spring break, summer vacation and winter vacation. In Singapore, elementary schools have four vacations or breaks: spring break, summer vacation, autumn break and winter vacation. For detailed timing of vacations or breaks of each region, please refer Figure 2.

**Figure 2.**
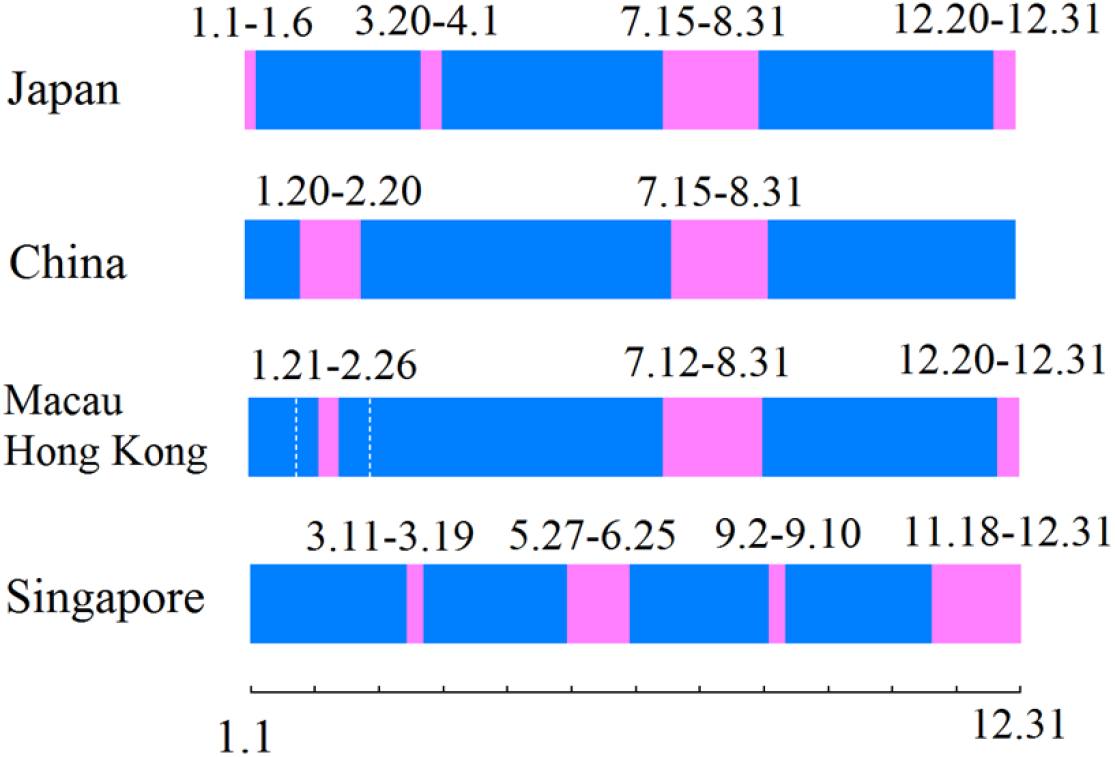
Time of school terms in mainland China, Hong Kong, Macau, Japan and Singapore. The dashed lines for Macau and Hong Kong show the time range that cover CSF from 2010 to 2017.

The time period of the SFTR in mainland China changes every year depending on the date of the CSF. Considering the earliest start time and the latest end time of the SFTR from 2009 to 2016, we used the time period from mid-January to mid-March as the SFTR period in mainland China.

### Climatic data

Climatic data for Hong SAR, Macau SAR and Japan were obtained from Hong Kong Observatory [29], Macau Meteorological and Geophysical Bureau [30] and Japan Meteorological Agency [31] respectively. Climatic data for the three provinces in China were obtained from the National Meteorological Information Center of China [32]. Climatic factors considered are average temperature, relative humidity, rainfall, and the hours of sunshine. There is no data for relative humidity in Japan.

Data collected are presented in Appendix 1 in the Supplemental Information.

## Methods

### Wavelet analysis of the periodicity of HFMD incidence

We used wavelet analysis method to analyze the periodicity of the time series of HFMD for selected countries, SARs and provinces in mainland China. To stabilize the variance, wavelet decompositions were performed to the square root of incidence data.

### MCMC-TSIR model to estimate the transmission rate

In order to analyze the seasonality of HFMD transmission rate, we applied the Time Series Susceptible Infected Recovered (TSIR) framework for the dynamic model of HFMD transmission [33]. The TSIR model has been applied to the analysis of transmission of childhood infectious diseases such as measles, rubella and chickenpox. Interested readers can refer to the relevant literature [9,34–36].

The number of infected in the model follows the negative binomial distribution *Y*(*t*+1)~*NB*(*λ*, *Y*(*t*)), where *λ* is the force of infection. The force of infection is modeled as *λ*=*β_m_Y^α^*(*t*)*X*(*t*), where *X*(*t*) and *Y*(*t*) are the number of susceptible and infected individuals respectively, *α*(0<*α*≤1) is the degree of population mixing, and *β_m_* is the month-specific transmission rate (or the 4-week-specific transmission rate). We assume that the transmission rate pattern is the same every year in a region, that is, the transmission rate for a month (or four-week) of the year is the same. When *α*=1, it means that the population is well mixed; when *α*<1, the population is not well mixed, that is, the contact rate between some people is higher than that between other people.

The incubation period of HFMD is 3 to 5 days, and the infectious period is from 7 to 10 days [37,38]. After the infected was recovered, they may still shed viruses for one or two weeks. Hence we choose the serial interval as one month for provinces in mainland China and Macau, and 4 weeks for Japan, Hong Kong and Singapore. Using the TSIR model, we estimate in each region 14 (or 15) parameters, including 12 (or 13) parameters for monthly (or four-week) transmission rate, the population mixing degree *α* and the initial number of susceptible.

The parameters in the TSIR model are estimated by the Markov Chain Monte Carlo (MCMC). The Markov Chain is generated by using Gibbs sampling. After 1000 initial iterations of burn-in, Markov chains were run 10 million iterations, and then the last 10,000 iterations sampled. In order to avoid the autocorrelation of the sampling results, the Markov chains were sampled every 100 iterations.

Since CDCs in Shandong, Hainan, and Guangdong introduced EV71 vaccination in the second half of 2016, the use of vaccines has little effect on the number of cases, so the effects of immunization will not be considered in the TSIR model here.

### Transmission rate seasonality and seasonal deviation

The seasonal deviation around the mean transmission rate and the seasonality The transmission rate seasonality was calculated as

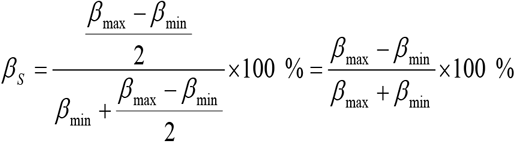

where *β*_max_ and *β*_min_ are the maximum and the minimum transmission rates. This is the same to that of the sinusoidal or the term time forcing [5].

We calculated the transmission rate seasonality using the above equation and we also calculated the seasonal deviation (in one-month interval) around the mean transmission rate:

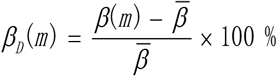

where, 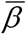 is the average of the transmission rate.

### The linear regression model of influential factors

In order to further analyze whether the HFMD transmission rate in each region is affected by climate factors and seasonal children’s contact, we use linear regression model for each region to analyze the association of transmission rate with climatic factors of monthly average temperature, rainfall, relative humidity and sunlight hours, and school terms. The SFTR period that represents the seasonal population contact rate will also be considered in the regression model for mainland China. Since the estimated transmission rate is the monthly value, we use February as the winter vacation time and July and August as the summer vacation time for provinces in mainland China. July and August are also the summer vacation time for Japan, Macau SAR and Hong Kong SAR. The annual SFTR time is not fixed; for years 2009 to 2016, the earliest start date is in the middle January and the latest ending date is in the middle of March. Considering that population flux may have delayed effect [36], we select February and March as the SFTR period with a two-week delay in the linear regression model.

## Results

### The periodicity of HFMD incidence in various regions

The HFMD incidence showed contrast periodicity that could be bi-ennual, annual or bi-annual periods in the countries or regions we studied (Figure 2). From the perspective of entire mainland China, the epidemic of HFMD had both annual and biannual dominant inter-epidemic periods. In the temperate zone, the dominant inter epidemic period of HFMD in Japan and Shandong was one year. Japan had an additional bi-ennual period from 2012 to 2016, and Shandong had an additional biannual inter-epidemic period from 2010 to 2012. In the subtropical zone, HFMD had a dominant annual inter-epidemic period in Hong Kong and Guangdong, but a bi-annual period after 2013 in Macau. HFMD in Singapore and Hainan that are both in the tropical zone had unstable inter-epidemic periods: in Singapore the dominant inter-epidemic period shifted from one-year to one-and-a-half-year before 2015 and then shifted to non-dominant period after 2015; in Hainan there was no dominant period before 2013, and mainly a one year period after 2013. We checked the dominant period for Singapore before 2009 and found that the period is also unstable (Appendix 2 in Supplemental Information). These results show that with different periodicities of inter-epidemic in a same climatic zone, the reason of forming the periodicity of HFMD incidence may be quite different.

**Figure 3.**
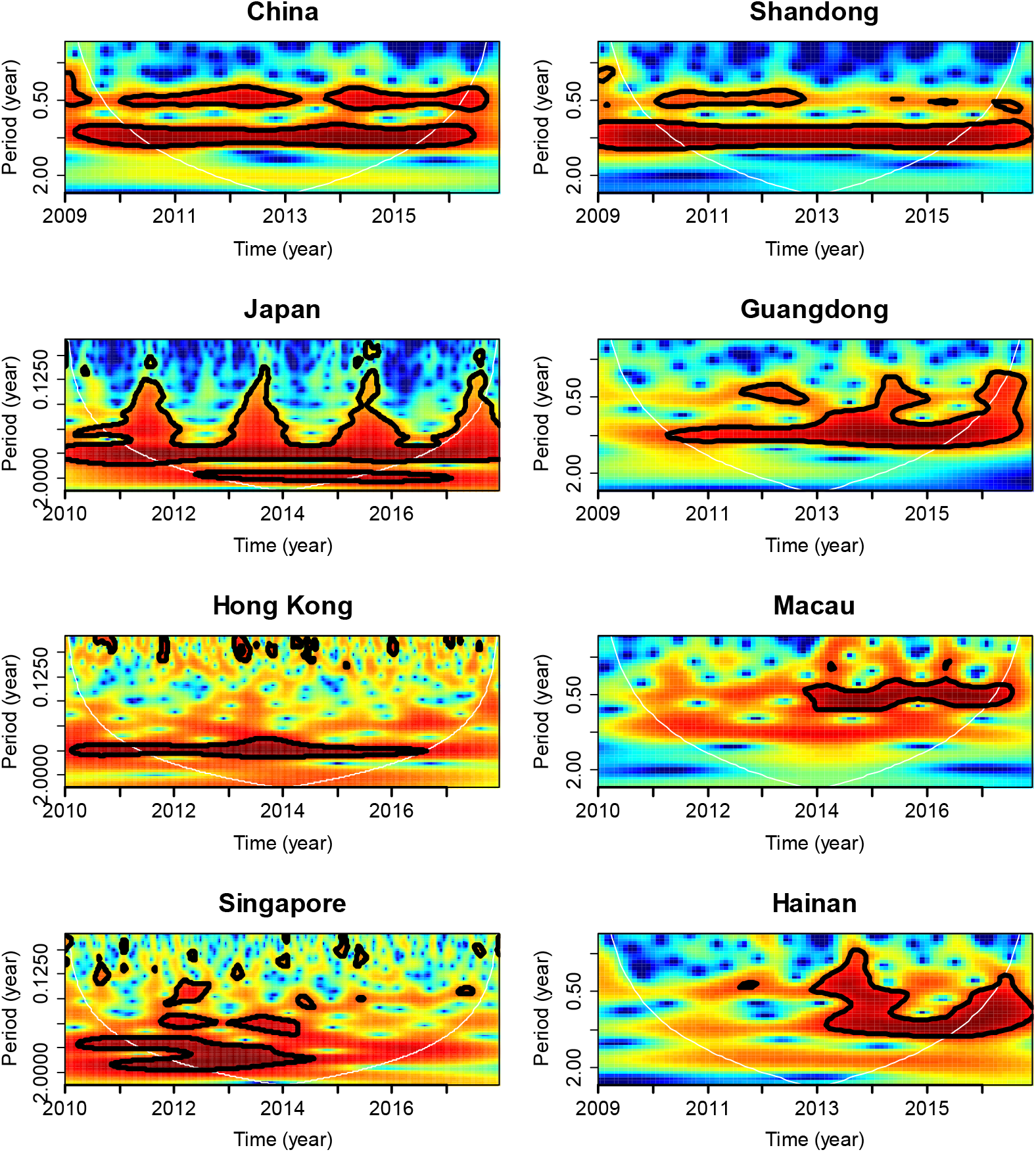
Results of wavelet analysis. Inter-epidemic periods in countries, SARs, and in provinces in mainland China.

### HFMD transmission rate and seasonality

The transmission rate seasonality of HFMD varied among countries, SARs and provinces in China (Figure 4). HFMD transmission in Singapore had no seasonality. In Japan, Macau SAR and Hong Kong SAR, the HFMD transmission rate were relatively higher in spring compared to other months of the year, however the peak month of transmission rate had a shift from April in Hong Kong and Macau to June in Japan. In provinces in mainland China and the entire country of China, the transmission rates of HFMD were all highly seasonal, and had a similar pattern: February to April were the peak months of HFMD transmission rate, and interestingly, there was a great increase of transmission rate in February. The increase in February was greater if we looked at the transmission rate at the country level, which supposed to be the average of transmission rate in all provinces in mainland China. The phenomenon of the dramatic increase of transmission rate in February was unique to mainland China and was not observed in Japan, Singapore, Hong Kong and Macau.

Estimated parameters are presented in Appendix 3 in the Supplemental Information.

**Figure 4.**
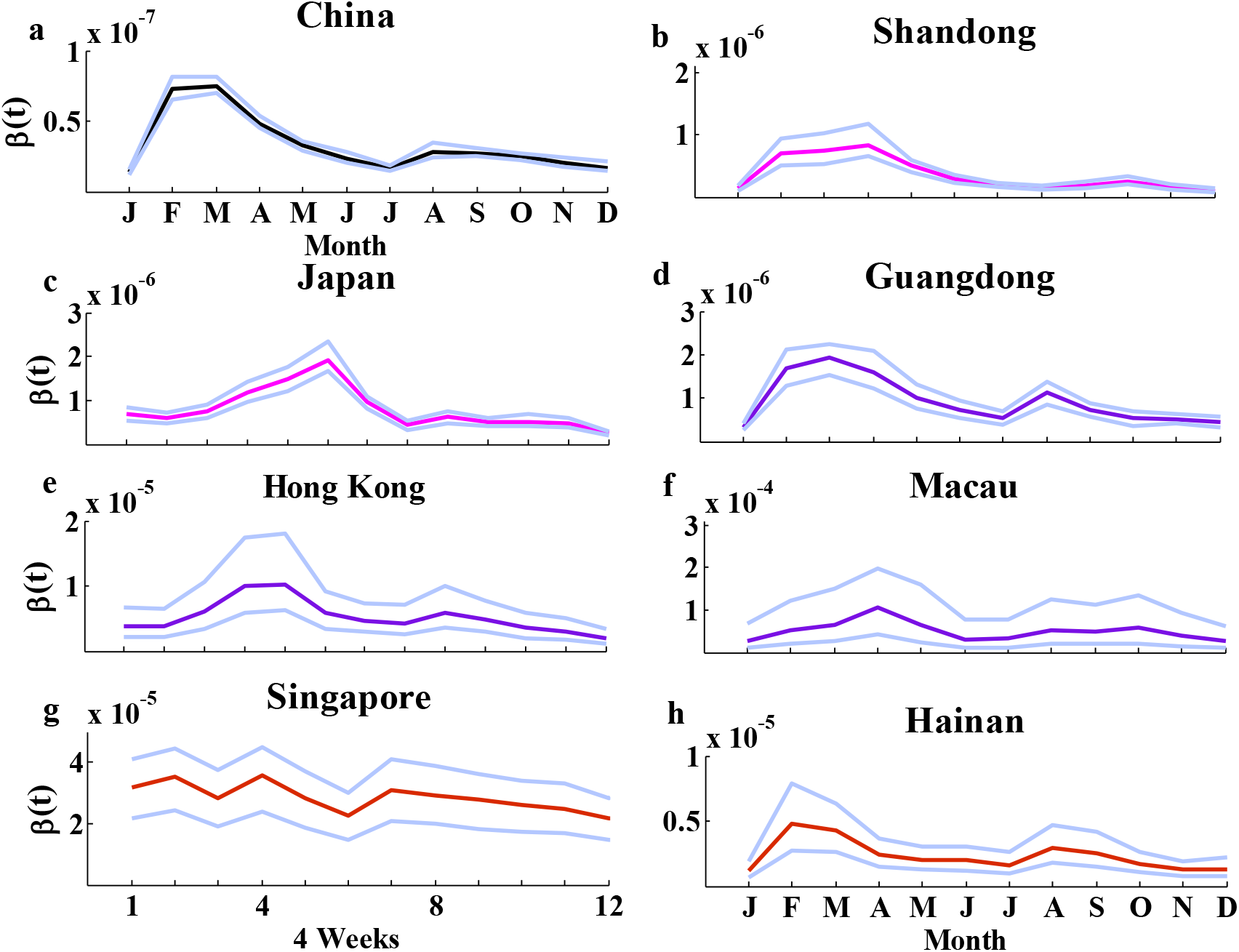
Estimated HFMD transmission rate with 95% Confidence Intervals. Colors of the transmission rate represent climatic zones: fuchsia, temperate zone; blue violet, subtropical zone; red, tropical zone. The upper and lower curves in light blue are the corresponding 95% confidence intervals.

### Possible reasons of HFMD transmission rate seasonality

The highest deviations around the mean transmission rate in regions except Singapore are similar, around 100% or higher (Figure 5). The transmission rate seasonality in these regions are also high, above 60% (Table 1).

**Figure 5.**
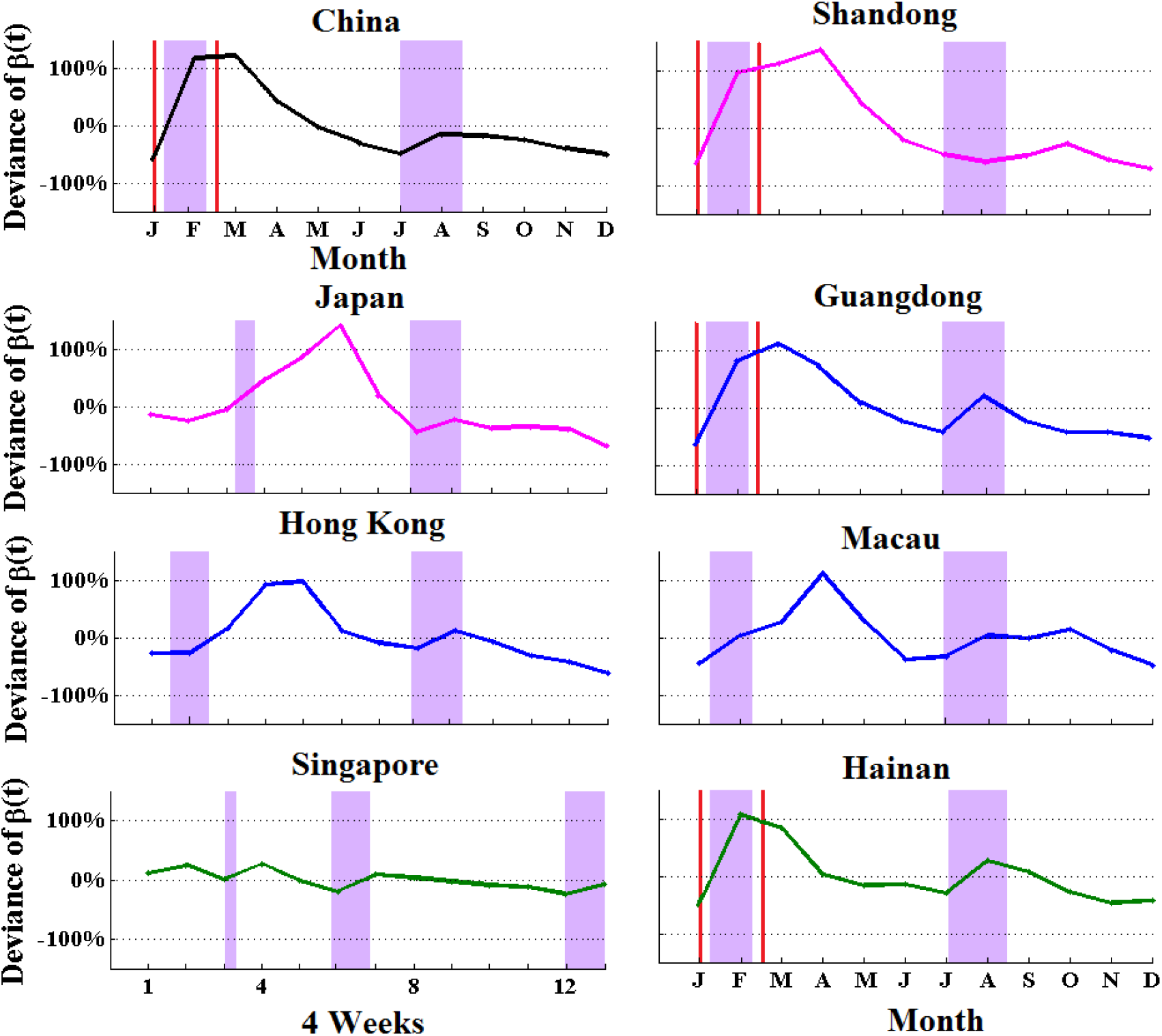
The deviation around the average transmission rate in each region, the pink shading shows the approximate timing of school vacations, and the red frame is the timing of SFTR.

**Table 1.** *R*_0_ and seasonality of transmission rate in countries, SARs and provinces in China

| | $R_0$ with 95% CI | Seasonality (%) |
| --- | --- | --- |
| Singapore | 53.1 [23.9, 95.7] | -- |
| Japan | 50.6 [41.5, 61.8] | 77.3 |
| Hong Kong | 18.9 [11.2, 31.9] | 67.8 |
| Macau | 15.1 [5.7, 34.5] | 60.9 |
| China | 22.6 [20.3, 25.7] | 69.1 |
| Hainan | 10.4 [6.2, 16.4] | 60.5 |
| Guangdong | 49.1 [37.5, 61.7] | 71.3 |
| Shandong | 17.3 [13.0, 22.7] | 78.1 |

School closure did not coincide with lower transmission rate for HFMD in all regions. This is quite different from the situation of measles or rubella in developed countries in the pre-vaccination era where during summer vacation the transmission rate was significantly lower than usual.

In the three provinces in mainland China and in the entire country, the transmission rate in February has increased significantly compared to the transmission rate in January. The large increase in the transmission rate within one month is unlikely to be caused by climatic factors, because no climatic factors have been observed to change rapidly in February. The dramatic increase in transmission rate in February coincides with the time period of SFTR, which depends on the time frame of CSF. Many things happen during this time period: 1) there is a large scale national population flux during SFTR that lasts 40 days and spans from mid-January to mid-March; 2) a large increase in the population contact rate during the CSF that is officially a one-week holiday; 3) Children in kindergartens or schools have a five-week winter vacation that is mainly during February. Combining these three situations the increase in the transmission rate in February is most likely due to the rapid increase in contact rates that is related to CSF.

### Regression results

Relative humidity in Hong Kong and Macau or rainfall in Japan has significant impact on HFMD transmission rate seasonality. However, in three provinces in mainland China, SFTR period has a significant effect on transmission rate, and in Guangdong both relative humidity and SFTR period have a significant effect.

**Table 2.** Model coefficients and statistics using regression models

| | Tempe<br>-rature | Relative<br>humidity | Rainfall | Sunshine<br>hours | School<br>Terms | SFTR<br>period | $R^2$ of<br>model |
| --- | --- | --- | --- | --- | --- | --- | --- |
| Japan | - | / | -5.925e-08* | - | - | / | 0.3235 |
| Hong Kong | - | 1.993e-7** | - | - | - | / | 0.5001 |
| Macau | - | 2.726e-6* | - | - | - | / | 0.3303 |
| Shandong | - | - | - | - | - | 4.485e-7* | 0.3578 |
| Guangdong | - | 5.476e-8* | - | - | - | 9.167e-7** | 0.7830 |
| Hainan | - | - | - | - | - | 2.655e-6*** | 0.7592 |

/ no data; \* $P < 0.1$ ; \*\* $P < 0.05$ ; \*\*\* $P < 0.01$ .

## Discussion

The incidence pattern of HFMD in different climatic regions could be quite different: biannual, annual, and occasional bi-ennual inter-epidemic periods. In regions in the temperate or subtropical zones, the inter-epidemic period was clear, but in Singapore that is in the tropical zone, the inter-epidemic period was unstable. This means that the epidemic of HFMD in Singapore was in chaos. Usually for childhood infectious diseases such as measles, high seasonality in transmission rate leads to chaos in epidemics, and low seasonality in transmission rate leads to annual seasonality in incidence [5,34]. Here we see a different situation. The transmission rate of HFMD is no seasonal in Singapore. However, the transmission rates of HFMD in some regions that have annual seasonality in incidence are obviously highly seasonal. How the transmission rate and its seasonality, interacted with birth rate, impact the incidence cyclic patterns of HFMD should be further studied.

The transmission rate of HFMD in three provinces in mainland China and the entire country has a similar seasonal pattern no matter that these provinces located in different temperate zones. Hence nationally there was a common driver for transmission rate seasonality for HFMD. The dramatic increase in transmission rate in February coincides with activities related to CSF. Activities are: SFTR that lasts for 40 days, winter school vacation that lasts one month or five weeks, and a week Spring Festival holiday. The population contact rate during the SFTR increased largely, but children’s contact rate with the same age group should be decreased during the winter vacation. Effects of these activities entangled and can not be distinguished in our study. The longest time period was SFTR, hence we used SFTR period to represent this special time period. We only studied the HFMD transmission rate including those during the time period of SFTR, but a more detailed study that can distinguish effects of activities should be conducted.

The population flux during SFTR is the highest in China and even in the world. In 2015, there were more than two billion travels during the SFTR. The migration population is mainly college students, peasant workers and their children, who go back and forth between their home towns and places they work or study at. This large-scale population migration in a short period of time makes the highest seasonal population contact rate in countryside and in large cities that was actually on the opposite. The population contact rate seasonality is a dominant factor for HFMD transmission in China.

On the contrary, the driver of transmission rate seasonality of HFMD in other countries or SARs is climatic factors. Yearly seasonal humidity or rainfall do affect the transmission rate of HFMD in these countries or regions. Singapore has a high level of humidity all year around. The less variation in key climatic factors such as relative humidity coincides with the non-seasonality in transmission rate in Singapore. Singapore, Hong Kong SAR and Macau SAR are listed as the highest population density country or territory in the world with population density of 8,274, 7,075 and 21,151 per square kilometer. But they also have high level of public health service and hygiene year around. These affect the transmission rate baseline but not seasonality. However, these regions do not have annual population contact rate seasonality. Compared with the four- to five-week winter vacation and SFTR in mainland China, Hong Kong and Macau have around 10 days holiday for Christmas at the end of December and one week for Spring Festival. Hence, the driving factor for transmission rate in these regions can be quite different than that in provinces in mainland China. The transmission rate of HFMD can be affected by the climatic factors as well as the seasonal contact rate, depending on which factor is dominant.

School terms did not significantly affect HFMD transmission rate. This is consistent with conclusions obtained from other studies for HFMD in Hong Kong and Singapore [39,40]: closing schools was found having a positive effect on reducing the spread of HFMD, but the impact was small. The average age of HFMD infection in China is around 3 years old, with a little younger age in provinces in the tropical and subtropical zones, and a little older in provinces in the north [2]. Most children, especially those in the southern part of China, may have contracted HFMD before going to kindergarten. Hence the effect of seasonal contact rate in children caused by school terms on HFMD transmission rate is not significant.

The impact of climatic factors on the transmission rate of viruses can be achieved by affecting the survival rate of the virus in the environment [41] or by changing social behavior [42]. When the survival rate of the virus in the environment is higher, more viruses will be accumulated in the environment, and indirectly increases the contact of susceptible individuals with the virus. On the other side, in raining, cold or hot days, people prune to stay indoor that could increase the contact rate. The HFMD is mainly fecal-oral transmitted and can be transmitted through contact with infected hands and contact with fomites. Hence not only washing hands is important to interrupt the transmission of HFMD but also cleaning of fomites is important.

Incidence cyclic pattern is shown to be determined by transmission rate and the number of susceptible [5,34]. There is no vaccination for HFMD, hence the susceptible pool is only changed by the birth rate. The birth rates in the studied provinces, SARs and countries are in a relatively small range of variation, the determinant of incidence pattern of HFMD should be the transmission rate baseline and seasonality. Because dominant factor for HFMD transmission seasonality in regions or countries other than mainland China is the climatic factors, the association of incidence and climatic factors is relatively consistent. However, in China, the dominant factor for HFMD transmission seasonality is seasonal population contact, hence we do not see consistent association of incidence and climatic factors.

## Acknowledgments

This work was supported by the Shandong Provincial Natural Science Foundation, China (Funding number: ZR2018MH037).

## Supporting Information Legends

Supplemental Information.pdf

